# Dynamic chromatin targeting of BRD4 stimulates cardiac fibroblast activation

**DOI:** 10.1101/563445

**Authors:** Matthew S. Stratton, Rushita A. Bagchi, Rachel A. Hirsch, Andrew S. Riching, Marina B. Felisbino, Blake Y. Enyart, Keith A. Koch, Maria A. Cavasin, Michael Alexanian, Kunhua Song, Jun Qi, Madeleine E. Lemieux, Maggie P.Y. Lam, Saptarsi M. Haldar, Charles Y. Lin, Timothy A. McKinsey

## Abstract

Small molecule inhibitors of the acetyl-histone binding protein BRD4 have been shown to block cardiac fibrosis in pre-clinical models of heart failure (HF). However, the mechanisms by which BRD4 promotes pathological myocardial fibrosis remain unclear. Here, we demonstrate that BRD4 functions as an effector of TGF-β signaling to stimulate conversion of quiescent cardiac fibroblasts into *Periostin* (*Postn*)-positive cells that express high levels of extracellular matrix. BRD4 undergoes stimulus-dependent, genome-wide redistribution in cardiac fibroblasts, becoming enriched on a subset of enhancers and super-enhancers, and leading to RNA polymerase II activation and expression of downstream target genes. Employing the SERTA domain-containing protein 4 (*Sertad4)* locus as a prototype, we demonstrate that dynamic chromatin targeting of BRD4 is controlled, in part, by p38 mitogen-activated protein kinase, and provide evidence of a novel function for *Sertad4* in TGF-β-mediated cardiac fibroblast activation. These findings define BRD4 as a central regulator of the pro-fibrotic cell state of cardiac fibroblasts, and establish a signaling circuit for epigenetic reprogramming in HF.

## Background

Fibrosis is a stereotypical wound-healing response that is mounted after tissue injury or stress. A significant body of clinical data has demonstrated that myocardial fibrosis is strongly associated with adverse outcomes in several forms of human heart failure (HF), including HF with reduced ejection fraction (HFrEF), HF with preserved ejection fraction (HFpEF), and genetically driven cardiomyopathies ^1,2^. While fibrotic responses may acutely serve to stabilize a focal area of myocardial damage, clinical and experimental studies support the contention that excessive, diffuse, or chronic activation of the fibrotic process can be deleterious to long-term cardiac function and patient survival. For example, interstitial fibrosis increases the passive stiffness of the myocardium, contributing to diastolic dysfunction ^3,4^, and disrupts electrical conduction in the heart, causing arrhythmias and sudden cardiac death ^5^. Unfortunately, despite the well-accepted roles of fibrosis in cardiac dysfunction, no targeted anti-fibrotic drugs for the heart exist. As such, understanding the fundamental mechanisms driving cardiac fibrosis and discovering novel approaches to target this process are of significant scientific and therapeutic interest.

The adult heart contains a large number of resident cardiac fibroblasts, which are developmentally derived from the epicardium and play important roles in maintaining tissue architecture ^6–8^. In response to injury, resident tissue fibroblasts undergo a dramatic cell state transition to become activated fibroblasts ^9^, sometimes referred to as myofibroblasts, which are characterized by expression of the marker gene *Periostin (Postn)* ^10,11^. This transition is associated with rewiring of the transcriptome to endow the activated fibroblasts with the capability to proliferate, migrate, secrete pro-inflammatory mediators and secrete fibrotic extracellular matrix (ECM). A causal role for activated fibroblasts in cardiac fibrosis was established employing mice in which a tamoxifen-inducible *Cre* cassette was used to selectively deplete *Postn*-positive fibroblasts using diphtheria toxin ^11^. In these mice, deletion of *Postn*-positive fibroblasts blunted cardiac fibrosis in response to angiotensin II infusion or myocardial infarction ^11^.

Cardiac fibroblast activation is driven, in part, by transforming growth factor-β (TGF-β), which signals through a heterodimeric cell surface receptor complex consisting of TGFβ receptor type I (TGFβRI) and II (TGFβRII) ^12,13^. Canonical TGF-β signaling leads to phosphorylation and nuclear import of SMAD transcription factors, which bind to regulatory elements in a variety of pro-fibrotic genes. Deletion of TGF-β receptors or the SMAD3 family member in *Postn*-positive fibroblasts suppresses cardiac fibrosis in response to pressure overload in mice ^14^. TGF-β can also promote fibrosis by stimulating SMAD-independent, non-canonical pathways, such as those governed by mitogen-activated protein kinases (MAP kinases). For example, cardiac fibroblasts lacking p38α are resistant to pro-fibrotic TGF-β signaling, and deletion of p38α in *Postn*-positive fibroblasts *in vivo* blunts cardiac fibrosis in response to myocardial infarction ^15,16^. Transcriptional effectors of non-canonical TGF-β signaling in fibroblasts include nuclear factor of activated T cells (NFAT), serum response factor (SRF) and myocardin-related transcriptional cofactors (MRTFs) ^17–21^.

Recently, a family of epigenetic reader molecules called bromodomain and extraterminal (BET) acetyl-lysine binding proteins was shown to control pathological fibrosis ^22^. JQ1, a first-in-class, potent and specific inhibitor of BET bromodomains that functions by competitively displacing BET proteins from acetylated-histones ^23^, was found to block cardiac fibrosis in mice subjected to pressure overload or myocardial infarction ^24–26^. Bulk RNA-seq analysis of left ventricular tissue homogenates demonstrated the ability of JQ1 to block expression of fibroblasts activation markers, including *Postn* ^25^. However, the mechanism by which BET protein inhibition suppressed fibrotic remodeling in the heart remained undefined. Here, we show that JQ1 potently inhibits cardiac fibroblast activation, and that the BET family member, BRD4, functions as a downstream effector that couples pro-fibrotic TGF-β signaling to the epigenome in cardiac fibroblasts. The findings reveal a critical role for a chromatin reader protein in the epigenetic control of pathological cardiac fibrosis and HF.

## Results

### BRD4 inhibition blocks TGF-β-mediated cardiac fibroblast activation

When placed in a collagen gel, activated fibroblasts exhibit the ability to contract due to expression of α-smooth muscle actin (α-SMA) ^27,28^, and this process can be enhanced by stimulation with TGF-β ^29^. To begin to address the potential role of BRD4 in the control of cardiac fibroblast activation, primary adult rat ventricular fibroblast (ARVF) cultures were incorporated into collagen gels and stimulated with TGF-β in the absence or presence of JQ1. JQ1 significantly reduced basal and TGF-β-mediated collagen gel contraction in association with a profound reduction in α-SMA protein expression (Fig. 1a-c). Using specific siRNAs, knockdown of BRD4, but not BRD2 or BRD3, suppressed TGF-β-mediated induction of α-SMA (Fig. 1d,e). These data suggest that BRD4 is the primary BET family member that regulates cardiac fibroblast activation.

**Figure 1.**
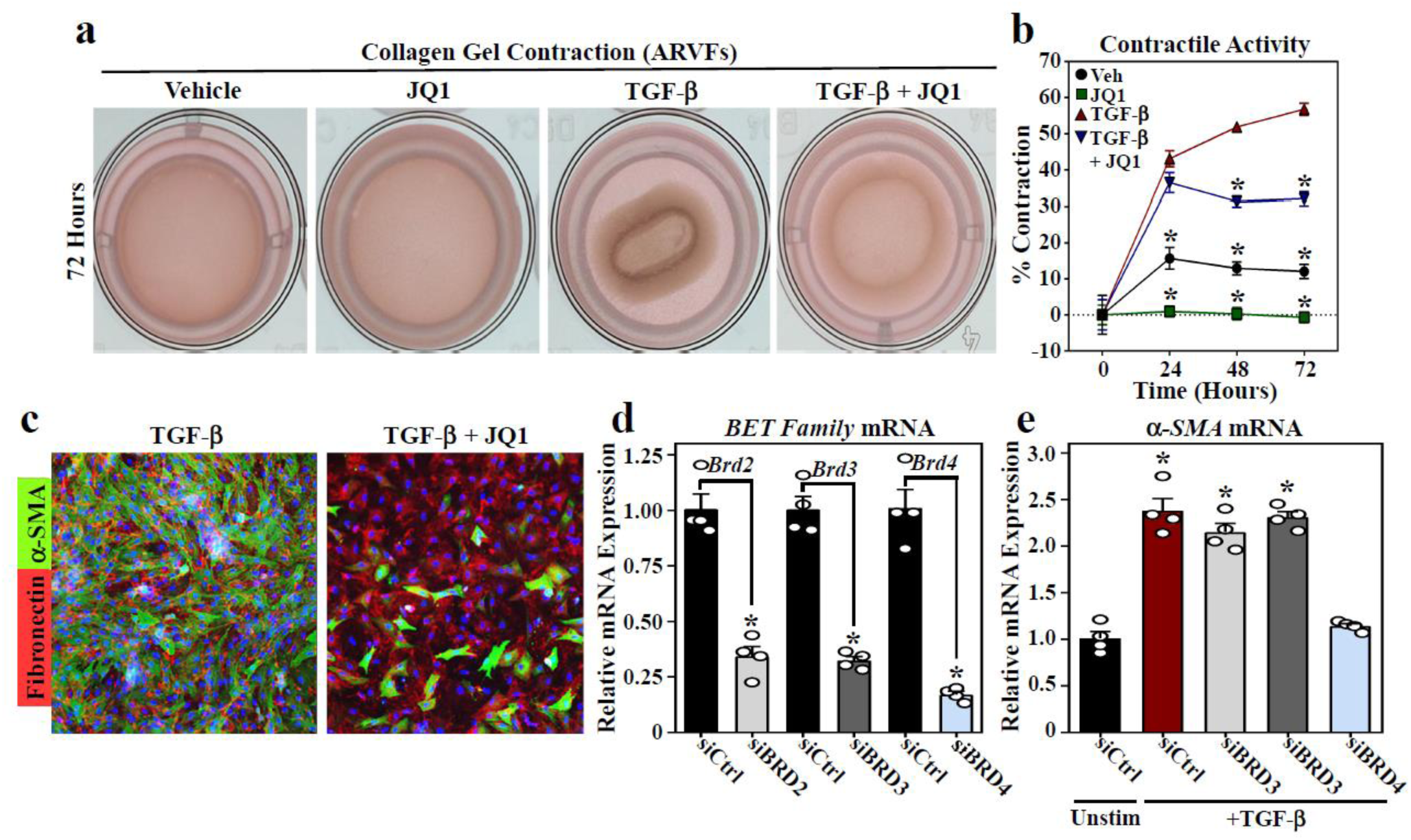
JQ1 suppresses TGF-β-induced cardiac fibroblast activation. (**a**) ARVFs seeded on compressible collagen gel matrices were assayed for gel contraction following treatment with TGF-β_1_ (10ng/mL) and/or JQ1 (500nM) for 72 hours. (**b**) Quantification of gel contraction images, reported as percentage contraction; (n=4 plates per condition). Data are presented as mean□±□SEM. \**P*<□0.05 by one-way ANOVA with Tukey’s post-hoc test. (**c**) ARVFs were treated with TGF-β_1_ in the absence or presence of JQ1 for 72 hours prior to fixation for indirect immunofluorescence detection of fibronectin (red) and α-SMA (green). Nuclei were stained with DAPI (blue). (**d**) ARVFs were transfected with negative control siRNA (siCtrl) or siRNAs targeting BRD2, BRD3 or BRD4 and maintained in low serum medium for 48 hours. BET family member mRNA expression was determined by qRT-PCR (n=4 plates of cells per condition). (**e**)ARVFs were transfected with the indicated siRNAs and maintained in low serum medium or were treated with TGF-β_1_ for 48 hours prior to determination of α-SMA mRNA expression levels by qRT-PCR (n=4 plates of cells per condition). All data are presented as mean□+SEM. Statistical analysis was performed by unpaired t-test (**d**) or one-way ANOVA with Tukey’s post-hoc test (**e**), \**P*□<□0.05 vs siCtrl.

### BRD4 inhibition blocks TGF-β-mediated mRNA and protein expression in cardiac fibroblasts

To further address the role of BRD4 in cardiac fibroblast activation, transcriptomic profiling by RNA sequencing (RNA-seq) was performed using RNA from ARVFs stimulated with TGF-β for 24 hours in the absence or presence of JQ1. Using a >two-fold expression change, minimum average expression of 5 FPKM, and a false discovery rate (FDR) of <0.05 as cutoffs, differential expression analysis showed that TGF-β increased expression of 174 genes and reduced expression of 236 genes which are depicted in heat map format (Fig. 2a). Data from cells treated with TGF-β in the presence of JQ1 are also included in the heat map. Ingenuity Pathway Analysis (IPA) of TGF-β-induced transcripts showed a strong enrichment for fibrosis and inflammatory signaling (Fig. 2b); the top 5 enriched pathways in TGF-β treated ARVFs in the absence or presence of JQ1 are shown. IPA also revealed that JQ1 profoundly suppressed expression of target genes that are known to be directly downstream of TGF-β signaling (Fig. 2c,d).

**Figure 2.**
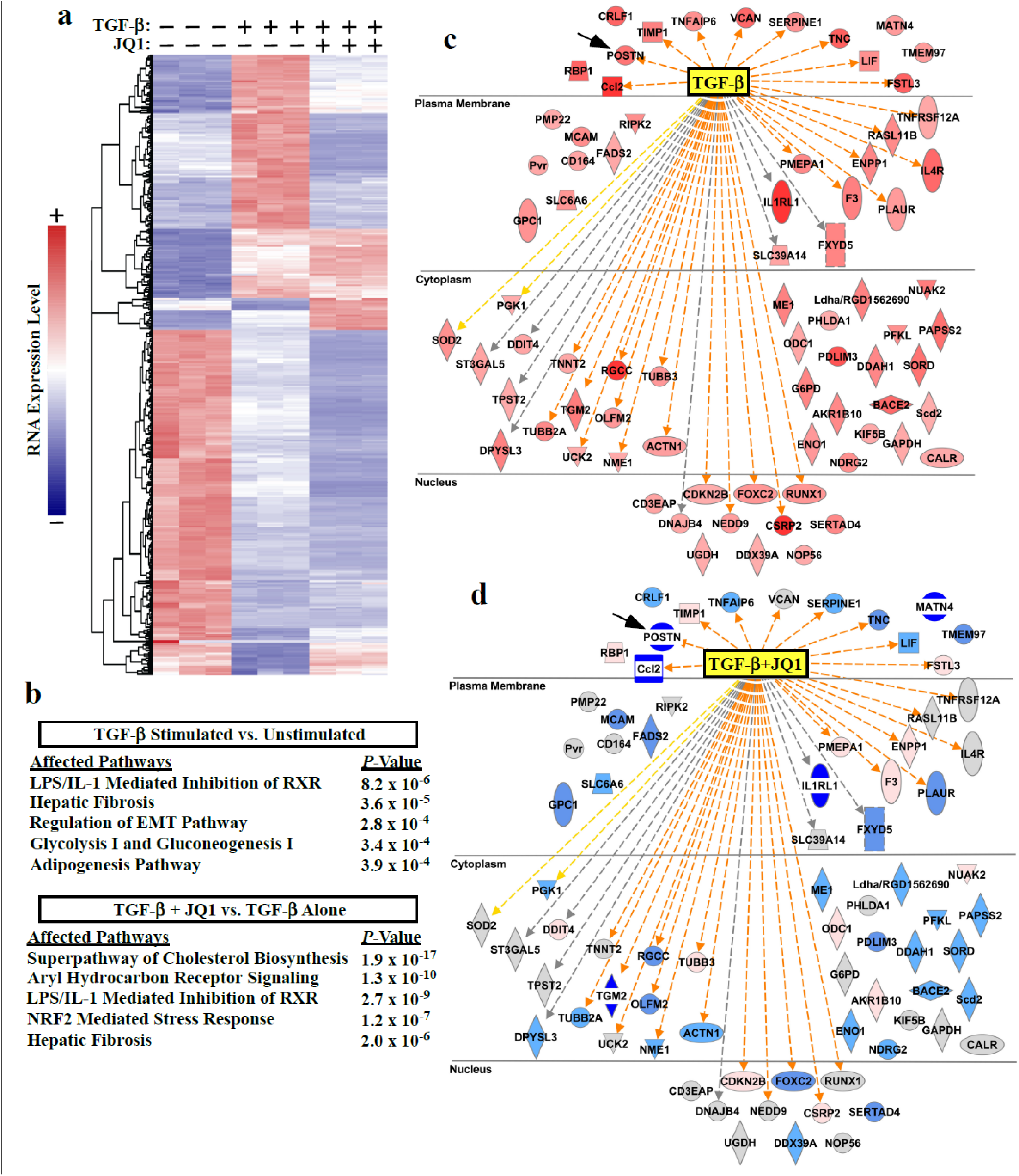
JQ1 globally suppresses pro-fibrotic gene expression in cardiac fibroblasts. ARVFs were treated with TGF-β_1_ in the absence or presence of JQ1 or vehicle control (DMSO) for 24 hours prior to extraction of RNA for RNA-seq analysis. (**a**) Heat map of significantly regulated genes. Each column represents data from a distinct plate of cells. (**b**) Ingenuity pathway analysis (IPA) was used to determine canonical pathways significantly altered by TGF-β stimulation relative to vehicle control or versus TGF-β + JQ1 treatment. The top five affected pathways for each analysis are reported; three redundant cholesterol biosynthesis pathways were removed from the lower table. (**c**) IPA analysis also indicated TGF-β as the strongest upstream effector molecule in the dataset. Induced genes that led to this determination are displayed (red indicates increased expression following TGF-β treatment). (**d**) Comparison of expression of these same genes in TGF-β-treated versus TGF-β + JQ1 treated cells revealed that JQ1 blocked induction of the majority of TGF-β downstream target genes (blue indicates decreased expression following JQ1 treatment).

*Postn* was among the most strongly downregulated transcripts in JQ1 treated ARVFs, further supporting a role for BRD4 in the control of cardiac fibroblast activation (Fig. 2c,d, black arrows). Quantitative mass spectrometry confirmed that the BET inhibitor blocked TGF-β-induced expression of Periostin protein (Supplemental Fig.1a). Furthermore, principal component analysis of relative protein abundance in whole-cell ARVF homogenates indicated clear segregations in the constellation of proteins expressed in unstimulated ARVFs compared to those in cells treated with TGF-β in the absence or presence of JQ1 (Supplemental Fig. 1b). Differential expression analysis revealed that TGF-β treatment significantly induced the expression of 64 proteins (adjusted *P*-value < 0.05). As depicted in the heat map, JQ1 blocked induction of a subset, but not all, TGF-β-induced proteins (Supplemental Fig. 1c; Periostin is indicated with a black arrow).

### BRD4 inhibition blocks TAC-induced changes in cardiac fibroblast gene expression *in vivo*

To determine whether BRD4 inhibition also suppresses cardiac fibroblast activation in the heart *in vivo*, mice were subjected to transverse aortic constriction (TAC) and treated with JQ1 or vehicle control by intraperitoneal injection every other day for two weeks (Fig. 3a). At the completion of the study, cardiac fibroblasts were isolated from the hearts of the mice as well as sham controls, and transcriptomic profiling of these cells was performed by RNA-seq; RNA was prepared from the cells immediately following isolation. Using a >two-fold expression change and a FDR <0.05 as cutoffs, 64 transcripts were found to be significantly upregulated in cardiac fibroblasts following TAC and significantly inhibited by JQ1 (Fig. 3b). Consistent with the cell culture data, TAC-induced *Postn* mRNA expression was dramatically repressed in fibroblasts from JQ1-treated mice (Fig. 3b, arrow). Additionally, IPA analysis revealed a strong enrichment for fibrosis and inflammatory signaling in cardiac fibroblasts post-TAC (Fig. 3c); the top 5 enriched pathways in cardiac fibroblasts from mice subjected to pressure overload in the absence or presence of JQ1 are shown. The impact of TAC and JQ1 on expression of fibrosis-and inflammation-associated mRNA transcripts is depicted (Fig. 3d). Together, the findings suggest that BRD4 regulates cardiac fibroblast activation in response to pressure overload *in vivo*.

**Figure 3.**
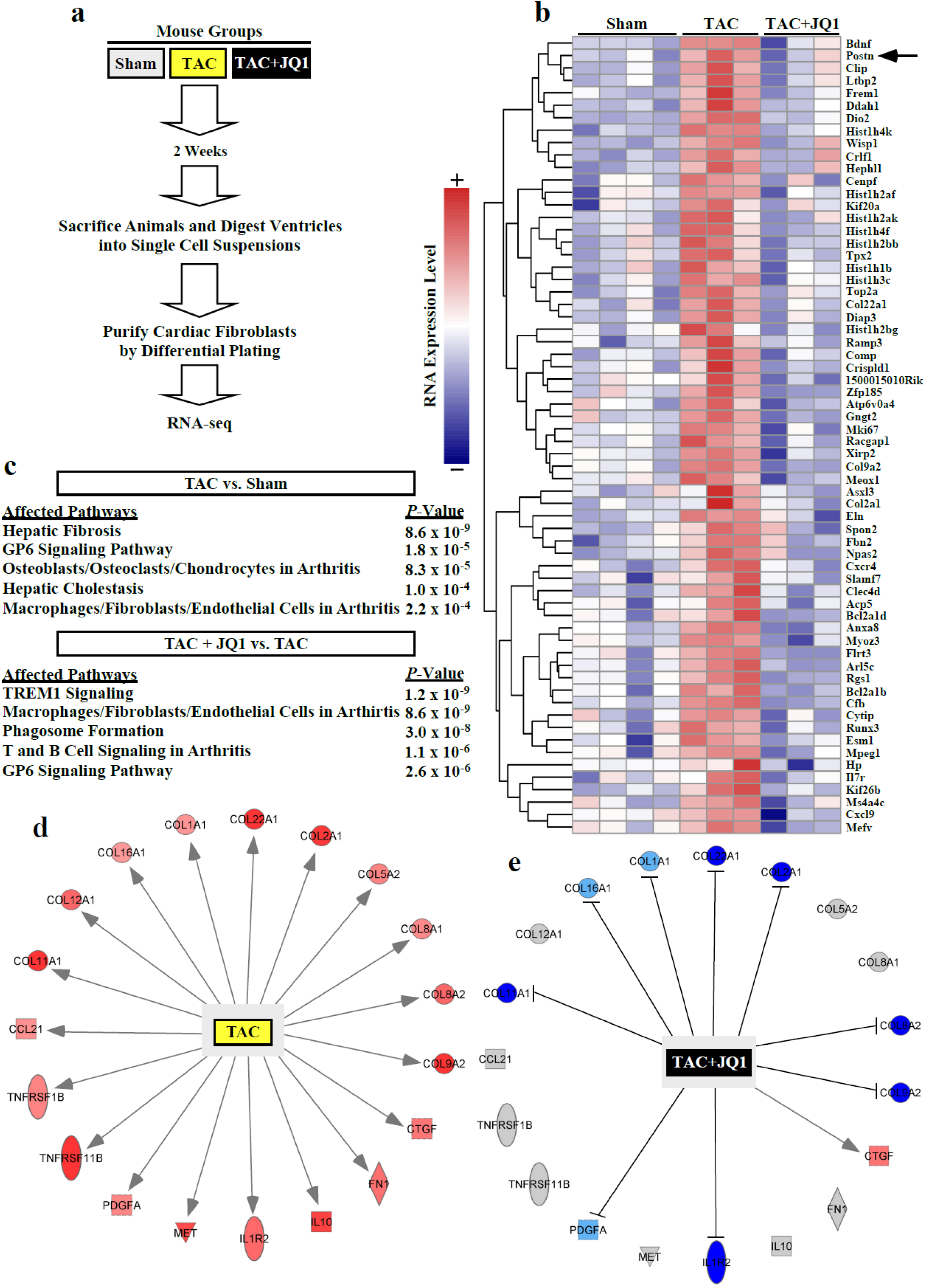
JQ1 suppresses pressure overload-induced pro-fibrotic gene expression in cardiac fibroblasts *in vivo*. (**a**) Schematic representation of the experiment. (**b**) Heat map of genes significantly upregulated by transverse aortic constriction (TAC) and suppressed by JQ1. (**c**) IPA was used to determine which gene expression pathways were significantly altered in cardiac fibroblasts isolated from mice subjected to TAC isolated versus Sham controls, or in cardiac fibroblasts from TAC +JQ1 treated mice versus TAC alone. The top five affected pathways for each analysis are shown. (**d**) The diagram depicts genes from the IPA ‘hepatic fibrosis pathway’ that were upregulated in cardiac fibroblasts from mice subjected to TAC (red indicates upregulation following TAC). (**e**) Comparison of expression of these same genes in TAC versus TAC + JQ1-treated mice revealed that JQ1 blocked induction of the majority of the target genes in this pathway (blue indicates decreased expression following JQ1 treatment).

### BRD4 binding to cardiac fibroblast enhancers is dynamically regulated by TGF-β

To address the mechanism by which BRD4 promotes cardiac fibroblast activation, ARVFs were subjected to whole-genome chromatin immunoprecipitation-sequencing (ChIP-seq) to map gene regulatory elements bound by BRD4. ARVFs were treated with vehicle control or stimulated with TGF-β for 24 hours prior to preparation of sheared chromatin (Fig. 4a). ChIP was performed separately with BRD4-and RNA Pol II-specific antibodies, and associated cardiac fibroblast DNA was subject to deep sequencing. Aggregate analysis of the two groups revealed well-defined peaks of BRD4 binding to 7,813 gene enhancers, with enhancers defined as being at least 2,500 base pairs (bp) from the transcription start site (TSS) of the associated gene (Fig. 4b). Prominent BRD4 binding was also detected at 4,069 gene promoters, defined as TSSs +/- 1000 bp (Supplemental Figure 2).

**Figure 4.**
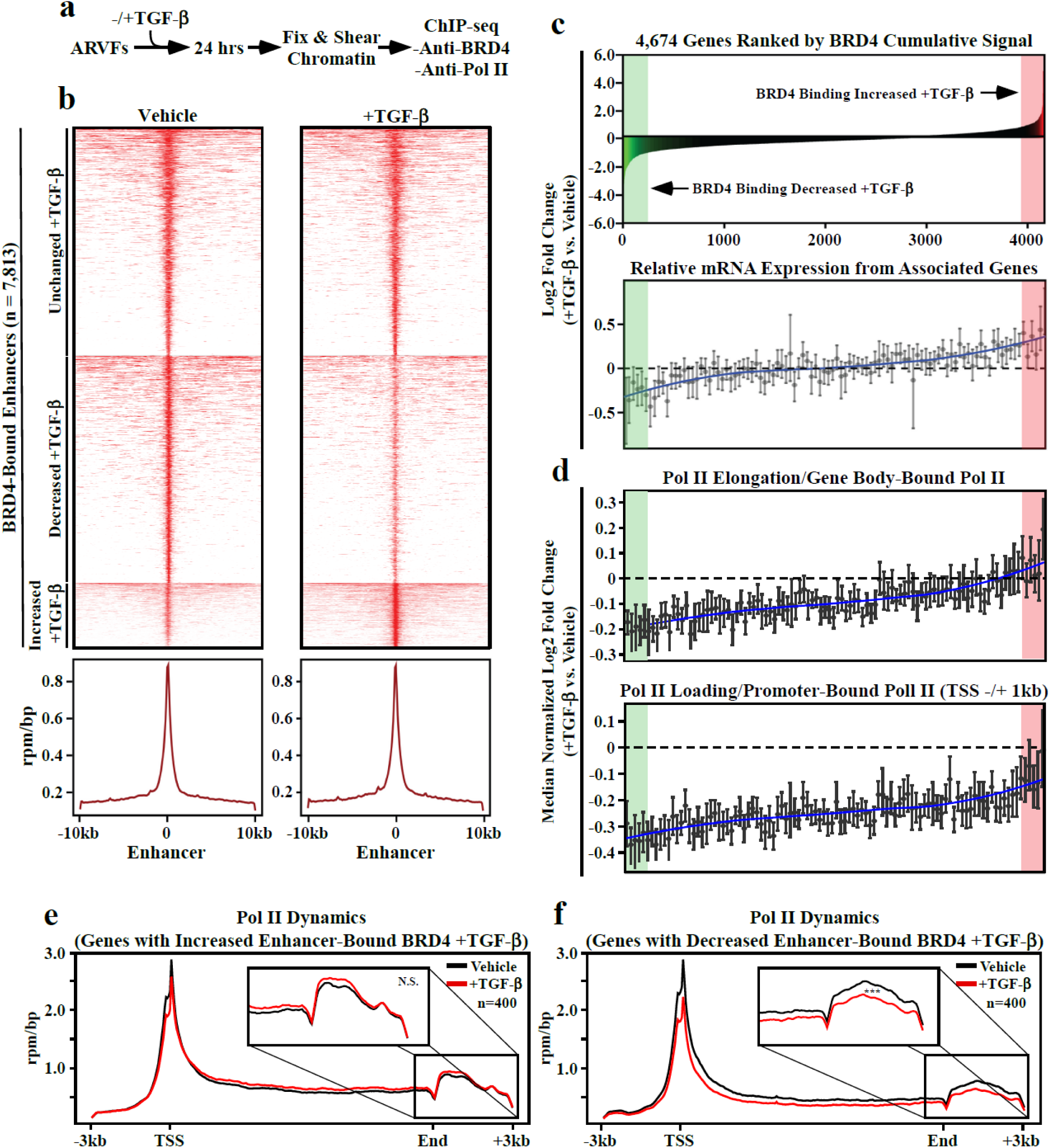
Chromatin targeting of BRD4 in cardiac fibroblasts correlates with RNA pol II elongation and downstream target gene expression. (**a**) Schematic representation of the experiment. (**b**) Heat map of BRD4-bound enhancers in TGF-β and unstimulated ARVFs covering 10kb upstream and downstream of the enhancer summit, where enhancers are grouped by increased, decreased or unchanged BRD4 binding in TGF-β stimulated ARVFs compared to unstimulated cells; mean reads per million mapped reads per base pair (RPM/bp). (**c**) Top – a histogram of cumulative BRD4 binding to enhancer/promoters of 4,674 active genes ranked by cumulative BRD4 enhancer abundance in response to TGF-β treatment. Bottom - a second histogram depicts the relative mRNA expression of genes proximal to the BRD4 bound enhancers/promoters. The downstream target gene mRNAs were clustered into bins of 100, ranked left-to-right based on cumulative BRD4 abundance at the enhancer/promoter (above); the data are presented as means ±SEM. (**d**) RNA Pol II binding to gene bodies (top) and promoters (bottom) of the corresponding genes is also shown as a histogram. RNA Pol II-bound genes were clustered into bins of 100, ranked left-to-right based on log2 fold-change of cumulative BRD4 abundance at the enhancer/promoter (above); the data are presented as means ±SEM. RNA Pol II rpm/bp plotted for 400 genes where TGF-β treatment led to increased BRD4 binding (**e**), or decreased BRD4 binding (*** denotes *P*-value < 10^9^) (**f**). The insets magnify the terminal regions of the genes, highlighting RNA Pol II elongation behavior.

To assess whether genomic targeting of BRD4 in cardiac fibroblasts is under signal-dependent control, we evaluated binding of the protein to 4,674 genes (enhancers and promoters combined), which are defined as active genes based on the presence of RNA Pol II in the TSS +/-1000 bp region, and an fpkm >= 10. Three general patterns of BRD4 dynamics were observed: (i) increased BRD4 binding upon TGF-β stimulation, (ii) loss of BRD4 binding following TGF-β stimulation, and (iii) constitutive BRD4 binding that is unchanged by TGF-β treatment (Fig. 4c, top panel); the 400 genes where BRD4 association with regulatory elements was most dramatically increased and decreased are highlighted in red and green, respectively. BRD4 abundance correlated with downstream target gene expression (Fig. 4c, bottom panel).

Comparison of BRD4 and RNA Pol II ChIP-seq signals suggested that association of BRD4 with regulatory elements for the 4,674 genes is coupled to productive transcription elongation, since RNA Pol II abundance within the bodies of the genes correlated with the degree of BRD4 binding (Fig. 4d, top panel); the amount of RNA Pol II within the gene body only increased in cases where TGF-β stimulation enhanced BRD4 binding. Paradoxically, TGF-β treatment led to decreased RNA Pol II binding to promoter elements for all of the 4,674 genes examined, although the degree of the diminution paralleled the extent of BRD4 binding (Fig. 4d, bottom panel).

RNA Pol II dynamics were further scrutinized by focusing on the top 400 genes where BRD4 binding was most dramatically increased or decreased in response to TGF-β. ChIP-seq showed that TGF-β stimulation enhanced gene-body enrichment of RNA Pol II when BRD4 was gained at enhancers/promoters, but decreased gene-body enrichment of RNA Pol II when the presence of BRD4 was reduced at regulatory sites upon cellular stimulation (Fig. 4e,f). These findings corroborate a critical role for BRD4 in TGF-β-coupled pause release of RNA poll II at upregulated genes in cardiac fibroblasts.

### TGF-β-mediated recruitment of BRD4 to *Sertad4* enhancers and super-enhancers

To further interrogate signal-dependent regulation of BRD4 genomic localization in cardiac fibroblasts, subsequent analyses focused on enhancers and super-enhancers (SEs), which are long-range gene regulatory elements that have been defined in cancer and immune cells based on abundant BRD4 binding above a threshold level found at typical enhancers ^30–32^. Dynamic, signal-regulated targeting of BRD4 in cardiac fibroblasts is exemplified by the *Sertad4* locus, where the chromatin reader was readily detected in association with the promoter region and six distinct proximal enhancers (E1 – E6) (Fig. 5a, top panel). TGF-β treatment led to enhanced binding of BRD4 to each of these sites, with profound enrichment occurring at E1 and E2 (Fig. 5a, bottom panel). Based on the level of BRD4 association, both E1 and E2 were categorized as SEs following stimulation with the pro-fibrotic agonist. In total, 147 and 116 BRD4-enriched SEs were detected in unstimulated and TGF-β stimulated cardiac fibroblasts, respectively (Fig. 5b). The ranking of E1 and E2, commensurate with BRD4 abundance before and after TGF-β stimulation, is indicated (Fib. 5b, yellow and green circles). Consistent with a regulatory role for BRD4, TGF-β-induced expression of *Sertad4* was blocked by JQ1 (Fig. 5c).

**Figure 5.**
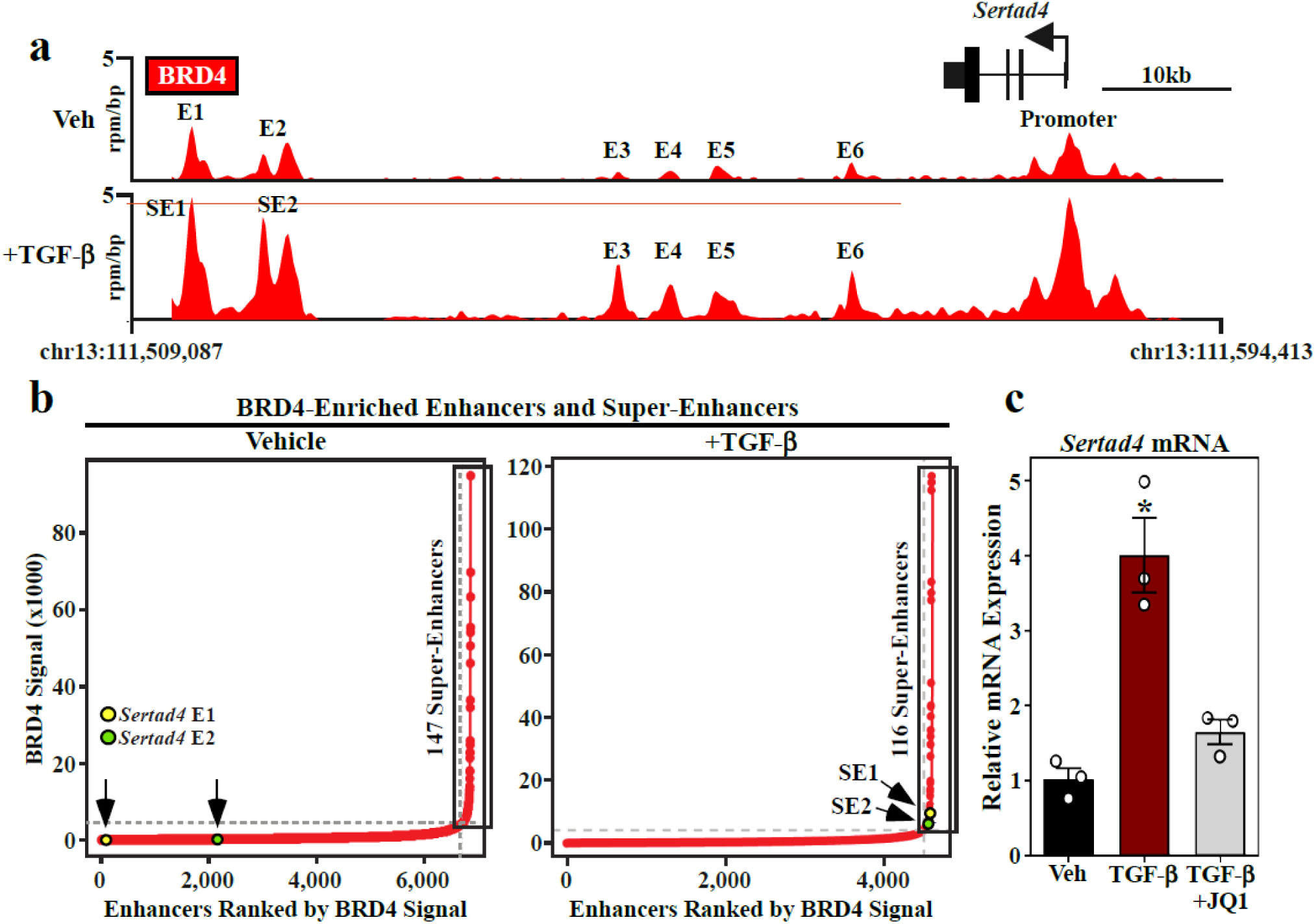
TGF-β stimulates binding of BRD4 to enhancers and super-enhancers associated with the *Sertad4* gene. (**a**) Top - gene track of the *Sertad4* locus showing BRD4 binding to the promoter and six distinct proximal enhancers (E) in unstimulated ARVFs. Bottom – upon TGF-β stimulation for 24 hours, BRD4 binding to all of these sites was increased, reaching a threshold for definition of E1 and E2 as super-enhancers (SE). (**b**) Hockey-stick plots of BRD4-enriched enhancers in unstimulated ARVFs and ARVFs stimulated with TGF-β for 24 hours. SEs, based on a threshold level of BRD4 binding, are boxed. The abundance of BRD4 at E/SE1 and E/SE2 of the *Sertad4* locus are indicated. (**c**) ARVFs were treated with TGF-β for 48 hours in the absence or presence of JQ1 or DMSO vehicle. Data are presented as mean□±□SEM. \**P*□<□0.05 vs vehicle by one-way ANOVA with Tukey’s post-hoc test.

### p38 MAP kinase regulates TGF-β-mediated recruitment of BRD4 to *Sertad4* enhancers and super-enhancers

The *Sertad4* locus was employed as a prototype to begin to define the signaling pathways that couple TGF-β receptor activation to chromatin targeting of BRD4 in cardiac fibroblasts. Initially, a panel of pharmacological inhibitors was employed to address whether known effectors of cardiac fibroblast activation regulate agonist-dependent expression of *Sertad4* (Fig. 6a). As shown in Figure 6b, inhibition of p38 mitogen-activated protein kinase, but not extracellular signal-regulated kinase (ERK), c-Jun N-terminal kinase (JNK) or the calcineurin phosphatase, suppressed TGF-β-induced *Sertad4* mRNA expression in ARVFs. Subsequently, ChIP-PCR was used to address the hypothesis that p38 kinase activity contributes to signal-dependent chromatin targeting of BRD4 (Fig. 6c). Consistent with the mRNA expression data, pan-p38 inhibition with SB203580 led to significant decreases in binding of BRD4 to the promoter region of the *Sertad4* gene, as well as each of the six enhancers/super-enhancers defined Figure 5. These data suggest that p38 signaling is involved in controlling BRD4 loading on gene regulatory elements in cardiac fibroblasts.

**Figure 6.**
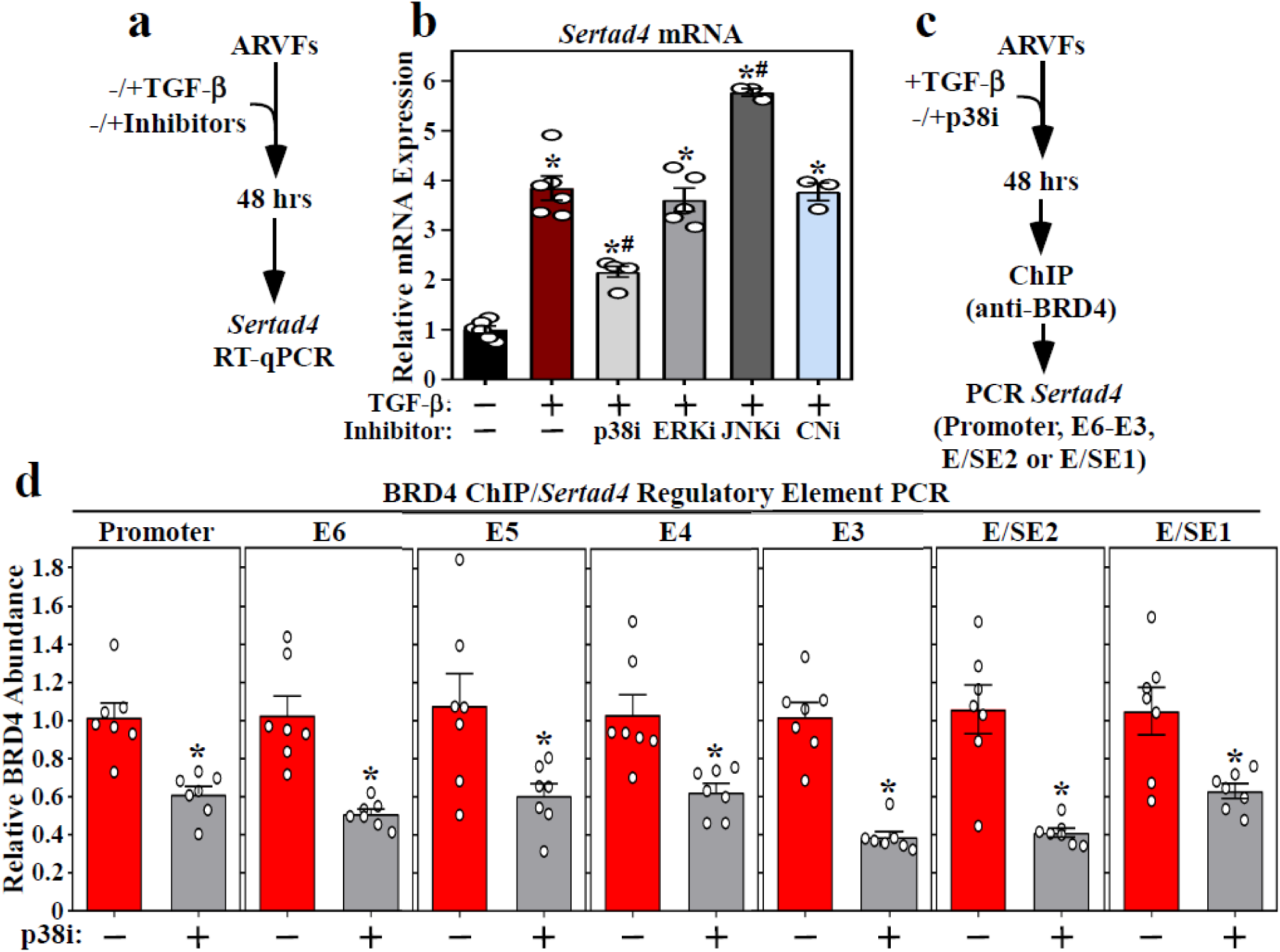
p38 inhibition suppresses recruitment of BRD4 to enhancers and super-enhancers associated with the *Sertad4* gene. (**a**) Schematic representation of mRNA expression experiment. (**b**) Inhibition of p38 (SB203580), but not ERK (PD98059), JNK (SP600125) and calcineurin (CN; cyclosporin A), blunted TGF-β-induced *Sertad4* Mrna expression in ARVFs. Data are presented as mean□+SEM; n=3-6 plates of cells per condition. \**P*□<□0.05 vs Vehicle, ^#^*P*□<□0.05 vs TGF-β alone by one-way ANOVA with Tukey’s post-hoc test. (**c**) Schematic representation of ChIP-PCR experiment. (**d**) BRD4-binding to the Sertad4 gene regulatory elements was significantly reduced in TGF-β stimulated ARVFs treated with the p38 inhibitor. Values were normalized to input levels are presented as mean□+SEM; n=7 per condition. \**P*□<□0.05 vs TGF-β alone by unpaired t-test.

### *Sertad4* regulates TGF-β-mediated induction of markers of cardiac fibroblast activation

*Sertad4* has not previously been implicated in the regulation of cardiac fibroblast activation. Thus, to address the functional relevance of BRD4-mediated control of this factor, RNA interference was used to knock down expression of *Sertad4* in ARVFs. Two independent lentivirus-encoded shRNAs efficiently repressed basal and TGF-β-induced expression of *Sertad4* (Fig. 7a). Furthermore, *Sertad4* knockdown markedly reduced TGF-β stimulated expression of *α-SMA* and *Postn* in the cardiac fibroblasts (Fig. 7b,c), suggesting that SERTAD4 protein contributes to cardiac fibroblast activation.

**Figure 7.**
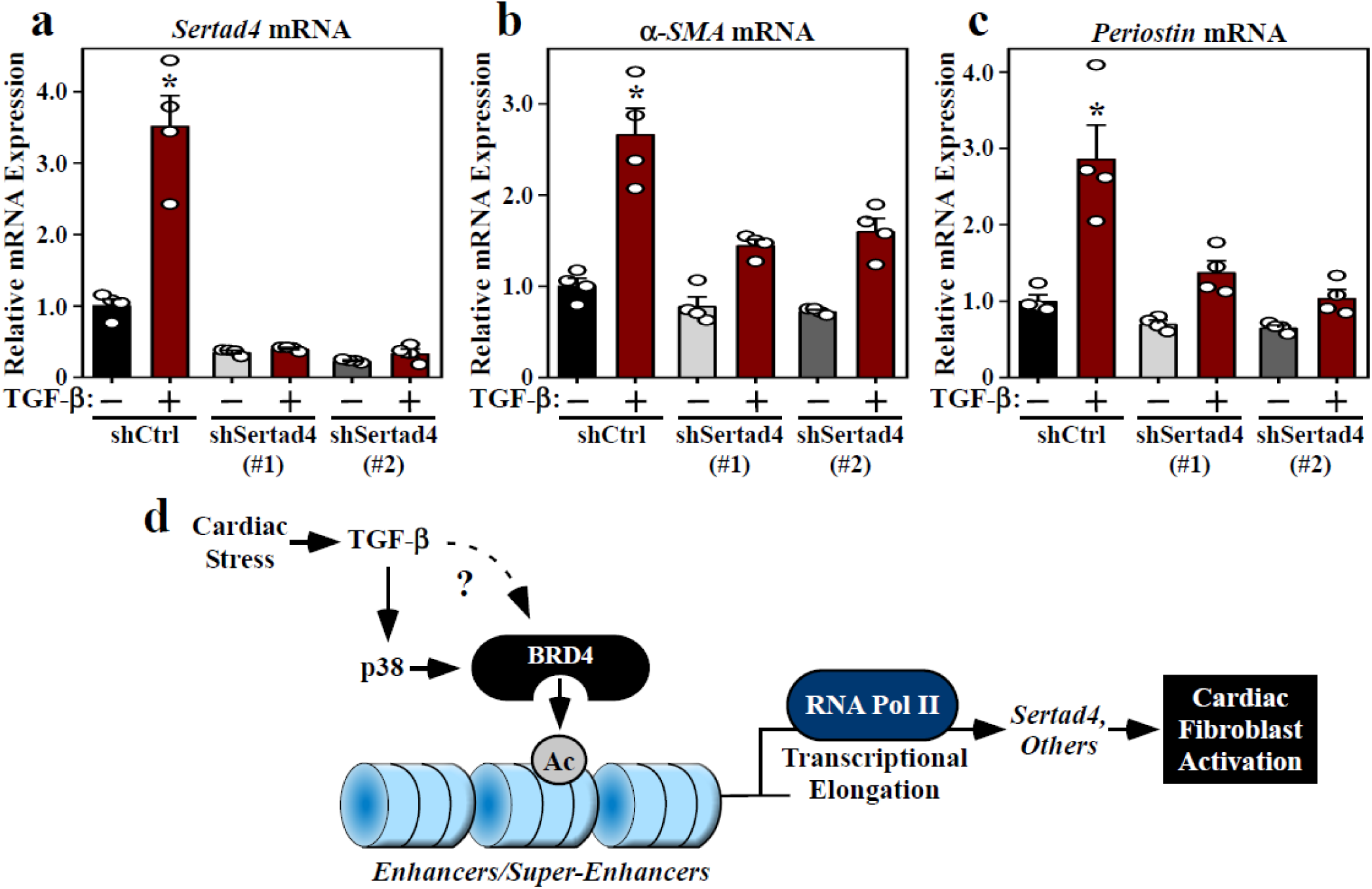
*Sertad4* knockdown represses cardiac fibroblast activation. ARVFs were infected with lentiviruses encoding control short-hairpin RNA (shRNA; shCtrl) or two independent shRNAs targeting Sertad4 (shSertad4 [#1] and [#2]. The cells were subsequently treated with TGF-β or DMSO control for 48 hours, and qRT-PCR analysis of *Sertad4* (**a**), α*-SMA* (**b**) and *periostin* (**c**) mRNA expression was performed. Data are presented as mean□+SEM; n=4 plates of cells per condition. \**P*□<□0.05 vs shControl by one-way ANOVA with Tukey’s post-hoc test. (**d**) A model for BRD4-mediated regulation of cardiac fibroblast activation. Cardiac stress signals, including TGF-β, stimulate p38 and likely other pathways to target BRD4 to gene regulatory elements, resulting in RNA Pol II elongation and expression of downstream targets, including *Sertad4*, which promote fibroblast activation.

## Discussion

Epigenetic mechanisms that regulate cardiac fibrosis remain poorly understood. Here, we demonstrate that the BRD4 chromatin reader protein is crucially involved in promoting cardiac fibroblast activation. BRD4 binding to discrete gene enhancers in fibroblasts can be altered by TGF-β receptor signaling in a p38 kinase-dependent manner, providing a circuit for coupling extracellular cues to the cardiac epigenome to drive pro-fibrotic gene expression (Fig. 7d).

Our prior RNA-seq analyses demonstrated the ability of JQ1 to block expression of canonical markers of fibroblast activation in the LV in mouse TAC and MI models ^25^. However, interpretation of these data was complicated by the use of bulk RNA from whole LV tissue homogenates. In the current study, we demonstrate that JQ1 profoundly inhibits expression of activation markers and ECM proteins in cultured cardiac fibroblasts treated with TGF-β, and in cardiac fibroblasts isolated from LVs of animals subjected to TAC. JQ1 also suppressed the contractile activity of cardiac fibroblasts in association with repression of expression of α-SMA. These data suggest that BRD4 functions in a fibroblast-autonomous manner to promote fibrotic remodeling of the heart.

We performed parallel ChIP-seq and RNA-seq to address, mechanistically, how BRD4 regulates fibroblast activation. Remarkably, to our knowledge, the resulting data represent the first genome-wide evaluation of epigenetic regulatory events in cardiac fibroblasts. The findings revealed that BRD4 is positioned at thousands of discrete enhancers and promoters throughout the cardiac fibroblast genome. In response to TGF-β treatment, the amount of BRD4 increases at a subset of these sites, while in other cases it decreases or is unaffected by the pro-fibrotic signal. This signal-dependent regulation of BRD4 is analogous to what we previously observed in cardiomyocytes treated with a hypertrophic agonist ^33^, although the genomic localization of BRD4 in fibroblasts versus myocytes is clearly distinct. In the current study, assessment of genes proximal to the BRD4 peaks demonstrated a correlation between BRD4 abundance at enhancers/promoters, RNA Pol II level within the gene body, and mRNA expression in cardiac fibroblasts. This relationship is consistent with the ability of BRD4 to stimulate gene expression by promoting transcriptional elongation and release of paused RNA Pol II ^34–36^. It is interesting to note that the number of BRD4-enriched enhancers in unstimulated cardiac fibroblasts actually decreased following TGF-β stimulation, suggesting that BRD4 activity needs to be repressed at certain genes in order for fibroblasts to become activated and secrete ECM. In other cellular contexts, coordinately deployed transcription factors recruit BRD4 to active *cis*-regulatory elements in a manner that is dependent on the underlying acetyl-histone landscape of chromatin ^37^. We posit that signal-dependent alterations in transcription factor occupancy and histone acetyltransferase/deacetylase activity govern genome-wide redistribution of BRD4 to drive cardiac fibroblast activation.

ChIP-PCR analysis demonstrated that treatment of cardiac fibroblasts with the p38 inhibitor, SB203580, blunted TGF-β-mediated recruitment of BRD4 to enhancers proximal to the gene encoding *Sertad4*, resulting in reduced *Sertad4* mRNA expression. p38 kinase has long been recognized as an effector of pathological cardiac remodeling ^38^, and a recent report demonstrated that p38α within cardiac fibroblasts mediates pro-fibrotic TGF-β signaling in the heart ^15,16^. Deletion of p38α (*Mapk14* gene) in cultured fibroblasts blocked differentiation of the cells into α-SMA-positive fibroblasts in response to TGF-β, angiotensin II (Ang II) or cyclic stretching. This block could be overcome by ectopic overexpression of the canonical transient receptor potential 6 (TRPC6) channel, a constitutively active form of the calcineurin phosphatase, or the serum response factor (SRF) transcription factor, suggesting that p38α lies upstream of these proteins, all of which have been demonstrated to regulate myofibroblast differentiation ^17^. Our data define BRD4 as another downstream effector of pro-fibrotic p38 signaling. The mechanism by which p38 controls BRD4, and the extent to which this kinase regulates the genome-wide distribution of BRD4 in cardiac fibroblasts and other cell types, remain to be determined.

SERTAD4 is a member of the SERTAD family of proteins ^39,40^, which are also known as Trip-Br or SEI proteins ^41^. SERTAD proteins are characterized by the presence of a conserved SERTAD (<u>SE</u>I, <u>R</u>BT1, <u>TA</u>RA) domain ^39,41,42^. Members of this family have been shown to regulate cell cycle progression by virtue of their ability to interact with CDK/cyclin complexes ^43^, and also by functioning as transcriptional co-factors to positively or negatively control E2F-dependent transcription ^42^. The biological function of SERTAD4, which is nuclear protein that appears to lack intrinsic transcriptional activity ^44^, is largely unknown. However, the related protein, SERTAD1, was found to act as a transcriptional co-activator of SMAD1 to stimulate BMP target gene expression ^45^. We demonstrate that TGF-β stimulates *Sertad4* expression, and that knockdown of this factor blunts TGF-β-induced expression of activation markers in cardiac fibroblasts. It is intriguing to speculate that, analogous to the BMP/SERTAD1 axis, SERTAD4 protein functions within a positive feedback circuit to promote TGF-β-dependent gene expression via SMADs, resulting in pathological cardiac fibrosis. This validation of SERTAD4 function highlights the ability of our epigenomic discovery approach to identify new regulators of cardiac fibroblast activation.

The BET family consists of BRD2, BRD3, BRD4 and BRDT, all of which contain tandem bromodomains. JQ1, which is a specific pan-inhibitor of all BET family members, has shape complementarity to the acetyl-lysine binding pocket within the bromodomains of these proteins ^23^. JQ1 competitively and reversibly displaces BETs from acetyl-histones on active enhancers, thereby disrupting downstream signaling events to RNA Pol II. We previously demonstrated that JQ1 treatment reduces interstitial fibrosis in mouse TAC and MI models ^24–26^. Recently, the ability of JQ1 to block cardiac fibrosis in the TAC model was confirmed by an independent group, and was associated with reduced endothelial to mesenchymal transition ^46^. JQ1 has also been shown to blunt fibrosis in other organ systems, including lung ^47–49^, kidney ^50^, liver and pancreas ^51,52^. Collectively, the data underscore the therapeutic potential of BET inhibitors for the treatment of diverse fibrotic diseases. Nonetheless, there is reasonable concern about potential toxicities of pan-BET inhibitors ^53^, which is highlighted by to adverse effects such as thrombocytopenia and nausea observed in clinical trials evaluating bromodomain inhibitors in cancer clinical trials ^54^.

BRD4 is a key pro-fibrotic BET family member. The data presented here enhance our understanding of the molecular mechanisms by which BRD4 controls pathogenic gene expression in fibroblasts. Together with forthcoming findings, this knowledge should yield a cumulative framework that guides the development of more selective BRD4 inhibitors, as opposed to pan-BET inhibitors, with the potential to provide a therapeutic window that is suitable for the treatment of chronic fibrotic diseases such as HF.

## Methods

### Cardiac fibroblast isolation and culture

Adult rat ventricular fibroblasts (ARVFs) were isolated from female Sprague Dawley rats using enzymatic digestion (1 mg/ml collagenase type II; Worthington Biochemical Corporation) followed by gentle centrifugation for collection of cells, as previously described ^55^. Cells were maintained in DMEM/F12 media supplemented with 20% FBS, 1 % antibiotics-L glutamine and 1 μmol/L ascorbic acid. ARVFs were then passaged once to P1 (passage 1) or twice to P2 (passage 2), and plated appropriately for downstream assays. Equilibration of cells in low serum medium (0.1% FBS) was carried out prior to all treatments. Cells were treated with the following agents (DMSO as vehicle; 0.1% final concentration): recombinant TGF-β_1_ (10ng/mL; Novoprotein), cyclosporin A (200nm; Sigma-Aldrich), JQ1 (500nM; synthesized in-house), SB203580 (10µM; Tocris), SP600125 (10µM; Selleckchem) or PD98059 (10µM; Selleckchem). For gene knockdown experiments using siRNAs, ARVFs were incubated with a transfection cocktail of siRNA (50nM; Sigma Aldrich) and Lipofectamine RNAiMAX reagent (Life Technologies) for eight hours, followed by removal of transfection complexes and treatment with TGF-β_1_ for an additional 48 hours. Lentiviral vector-mediated gene short-hairpin RNA (shRNA) knockdown experiments were performed by incubating ARVFs with lentivirus culture supernatants overnight, followed by equilibration with low-serum medium for an additional 24 hours prior to treatment with TGF-β_1_. Cells were harvested for RNA, protein or chromatin analysis as appropriate. Adult mouse cardiac fibroblasts were also isolated from murine hearts post-TAC (± JQ1 *in vivo* treatment) treatment using enzymatic tissue dissociation (Liberase TH Research Grade; Roche), as described previously and with minor modifications ^55,56^. Cells were pre-plated for one hour followed by washing and isolation of total RNA and processed for RNA-seq.

### Plasmids and lentivirus production

pLKO.1 plasmids (Sigma) encoding shRNA for *Sertad4* (TRCN0000247967 and TRCN0000247969) and a negative control (SHC002) were obtained through the Functional Genomics Facility at the University of Colorado Cancer Center. Lentiviruses were generated by co-transfecting L293 cells with pLKO.1 vectors in combination with packaging plasmid (psPAX2) and envelope plasmid (pMD2.G). Virus-containing cell culture supernatants were collected, filtered and used for experiments. psPAX2 and pMD2.G were gifts from Didier Trono (Addgene).

### Collagen gel contraction assay

Compressible collagen matrices were prepared using PureCol EZ gel solution (5mg/mL; Advanced BioMatrix). ARVFs suspended in serum-supplemented DMEM/F-12 medium were seeded (5□×□10^4^ cells/well) on collagen gels for 24 hours prior to equilibration by serum deprivation overnight. At the initiation of contraction, gels were released from wells and treated with DMSO (vehicle; 0.1% final concentration), TGF-β_1_ (10ng/mL) and/or JQ1 (500nM) for 72 hours. Well images were captured at 0, 24, 48 and 72 hours. Gel area for each well was determined using NIH ImageJ software and reported as percentage contraction.

### Mouse transverse aortic constriction (TAC) model

All animal experiments were performed in accordance with the Institutional Animal Care and Use Committee at the University of Colorado Denver. Ten week-old male C57BL/6 mice (Jackson Laboratories) were subjected to sham or TAC surgery using a 27-gauge needle to guide suture constriction ^26^. JQ1 was delivered every other day via intraperitoneal injection at a concentration of 50mg/kg in a 1:4 DMSO:10% (2-Hydroxypropyl)-β-cyclodextrin (Sigma Aldrich; 389145) vehicle starting the day of TAC surgery. Treatment groups were weight-matched prior to initiating the study.

### Mass spectrometry

Total protein was harvested from cardiac fibroblasts (three biological replicates) treated with DMSO (vehicle; 0.1% final concentration), TGF-β_1_ (10ng/mL) and/or JQ1 (500nM) using Mammalian Protein Extraction Reagent (M-PER; Thermo Scientific) and sonication (Fisher Scientific FB120). Protein concentration was determined using a bicinchoninic acid assay (Thermo Scientific). The samples (50µg) were digested using a filter-assisted sample preparation (FASP)/trypsin protocol. The digested peptides were fractionated using a high-pH reversed phase (RP) peptide fractionation kit (Thermo Scientific). Fractions collected from the high-pH RP fractionation were further separated using on-line second-dimension low-pH reversed-phase chromatography via an Easy-nLC 1200 nanoflow-UHPLC system (Thermo Scientific) on an EasySpray C18 column (PepMap RSLC C18, 3-μm particle, 100-Å pore; 75 μm x 15 cm dimension; Thermo Scientific) held at 45°C. The typical solvent gradient was 0–80 min: 0–35% B; 80–85 min: 35–100% B; 85–90 min: 100% B; 300 nL·min-1; Solvent A: 0.1% formic acid in water; Solvent B: 80% acetonitrile, 0.1% formic acid in water. 4 μL of each high-pH fraction was injected by the autosampler on the Easy-nLC 1200 nanoflow-UHPLC system. The eluent from the Easy-nLC system was analyzed using a Q-Exactive HF Hybrid Quadrupole-Orbitrap mass spectrometer (Thermo Scientific) coupled online to the nanoflow-UHPLC through a Thermo EasySpray ion source. Acquired mass spectra were centroids and converted to .mzML formats using ProteoWizard msConvert ^57^. Database search was performed using a modified SEQUEST algorithm implemented in Comet (Crux version 3.0 distribution) ^58^ with a concatenated decoy search against a *Rattus norvegicus* proteome database (TrEMBL and Swiss-Prot) retrieved on 2017/10/19 with 29,979 entries from http://uniprot.org ^59^. False discovery rate (FDR) analyses were performed by Percolator (Crux version 3.0) ^60^, requiring < 1% global peptide false discovery rate (Percolator q < 0.01) for a peptide and protein to be identified and quantified. Normalized spectral abundance factor (NSAF) values (calculated using peptides with Percolator q < 0.01) were calculated for protein quantification. Comparisons between the sample groups were performed using limma ^61^ in R/Bioconductor ^62^ with a moderated t-test using NSAF values. Technical replicate (two each) values were averaged prior to limma analysis. Proteomics data used for principal component analysis (PCA) were preprocessed by dividing protein NSAF values by the average of the three vehicle biological replicates. Proteins from the treatment groups vehicle, TGF-β, and TGF-β +JQ1 that were shown to be induced by TGF-β (Benjamini & Hochberg adjusted *P*-value ≤ 0.1) were extracted, log_2_ transformed, clustered, and plotted as a heat map using R.

### Quantitative real-time PCR (qRT-PCR)

Total RNA was collected from cells using QIAzol lysis reagent (Qiagen) and used for cDNA synthesis (Verso cDNA Synthesis Kit; Life Technologies). Gene expression was assayed on a StepOnePlus Real-Time PCR (Applied Biosystems) using the PowerUp SYBR Green Master Mix (Applied Biosystems). Amplicon abundance was quantified using the 2^-ΔΔCt^ method. Primer sequences are listed in Table S1.

### RNA-seq analysis using freshly isolated mouse cardiac fibroblasts

Total RNA was prepared from fibroblasts isolated from mouse hearts using Direct-zol RNA miniprep kit (Zymo Research). cDNA libraries were prepared using the NEBNext Ultra Directional Library Preparation Kit following rRNA depletion using the NEB rRNA depletion kit. Libraries were submitted to the UC Denver Genomics and Microarray core facility for sequencing (HiSeq2000). Reads were trimmed with Trim Galore! (v.0.3.7) and aligned with STAR (v.2.5.2a) ^63^ to the mouse genome (Mm10) with annotations from UCSC (July 2016). Raw counts were loaded in to R ^64^. Genes with a median of <1 mapped read per million were removed before differential expression analysis with edgeR ^65^. Raw and processed RNA-seq data were deposited to the GEO online database (http://www.ncbi.nlm.nih.gov/geo/) under accession number GSExxxxx.

### ChIP-seq and ChIP-PCR

Anti-BRD4 (301-985, Bethyl Laboratories) and anti-RNA Pol II (39097, Active Motif) Chromatin Immunoprecipitation was performed as previously described ^33^. Five independent 10-cm plates of ARVFs were combined per immunoprecipitation. Libraries were prepared using the Rubicon TruPLEX Tag-seq kit and submitted to the UC Denver Genomics and Microarray core facility for sequencing (HiSeq2000). For validation of BRD4-bound enhancers or super-enhancers of the *Sertad4* locus, ChIP was performed as described above. The immunoprecipitates were collected and eluted DNA was subjected to qPCR using specific primers (Integrated DNA Technologies; primer sequences are listed in Table S2.

### RNA-seq and analysis using ARVFs

ARVFs were treated with DMSO (vehicle; 0.1% final concentration), TGF-β_1_ (10ng/mL) and/or JQ1 (500nM). Total RNA was isolated using the Direct-zol RNA miniprep kit (Zymo Research). cDNA libraries were prepared using the NEBNext Ultra Directional RNA Library Prep Kit for Illumina following PolyA selection. Samples were submitted to WIGTC for sequencing (HiSeq 2500). All analyses were performed using the Rat RN6 genome and RN6 RefSeq gene annotations. Raw and processed ChIP-seq and RNA-seq data were deposited to the GEO online database (http://www.ncbi.nlm.nih.gov/geo/) under accession numbers GEO: GSExxxxx and GSExxxxx. All RNA-seq datasets were aligned to the transcriptome using Tophat2 (version 2.0.11) (http://www.genomebiology.com/2013/14/4/R36/abstract). Gene expression values were quantified using Cufflinks and Cuffnorm and differential expression was determined with Cuffdiff (version 2.2.0) ^66^.

### ChIP-seq analysis

All ChIP-seq datasets were aligned using Bowtie2 (version 2.2.1) to build version RN4 of the rat genome ^67^. Alignments were performed using the following criteria: -k 1. These criteria preserved only reads that mapped uniquely to the genome. ChIP-seq read densities were calculated and normalized using Bamliquidator (https://github.com/BradnerLab/pipeline/wiki/Bamliquidator). Briefly, ChIP-seq reads aligning to the region were extended by 200 base pairs (bp) and the density of reads per bp was calculated. The density of reads in each region was normalized to the total number of million mapped reads producing read density in units of reads per million mapped reads per bp (rpm/bp). MACS version 1.4.2 (model-based analysis of ChIP-seq) peak finding algorithm was used to identify regions of ChIP-seq enrichment over background ^68^. A *P*-value threshold of enrichment of 1e-9 was used for all datasets. A gene was defined as actively transcribed if enriched regions for RNA Pol II were located within 1 kb in either direction of the TSS.

### Mapping and comparing enhancers and super-enhancers

BRD4 ChIP-seq data were used to identify active *cis*-regulatory elements in the genome. ROSE2 (https://github.com/bradnerlab/pipeline/) was used to identify BRD4 enhancers and super-enhancers, as previously described ^30^. Briefly, proximal regions of BRD4 enrichment were stitched together if within 2 kb of one another. This 2 kb stitching parameter was determined by ROSE2 as the distance that optimally consolidated the number of discreet enriched regions in the genome while maintaining the largest fraction of enriched bases per region. Comparison of BRD4 changes at enhancers, super-enhancers, and promoter regions was performed as previously described ^30^. Active genes, defined by presence of RNA Pol II in the TSS +/-1 kb region, within 50 kb of enhancer regions were assigned as target genes, as described previously ^30^. Promoters for heat maps were identified by overlapping TSS -/+ 1,000 bp regions with peaks from treated and unstimulated BRD4 ChiP-seq data. Enhancer and promoter heat map figures were generated by combining three separate heat maps made from partitions of all the enhancers identified in the unstimulated and stimulated samples and all of the promoters identified in the unstimulated and stimulated samples. The partitions consisted of conserved regions (appearing in both states), lost regions (appearing only in the unstimulated state), and gained regions (appearing only in the stimulated state).

### Indirect immunofluorescence

ARVFs were cultured on 96-well clear-bottom plates (Greiner); each well received 3,500 cells in 100 μL of medium. The following day, cells were washed with serum-free medium and maintained in high glucose (4.5g/L) DMEM supplemented with penicillin-streptomycin. After incubation with DMSO or JQ1 for 1 hour, TGF-β_1_ (2ng/mL, Preprotech) was added to cells for 72 hours prior to fixation with 100% methanol (Fisher) at −20 °C. Fixed cells were washed and incubated with primary antibodies as a cocktail (1:2000 anti-α-SMA, Abcam, ab7817; 1:1000 anti-fibronectin, Abcam, ab2413) overnight at 4°C. Cells were washed and incubated with a cocktail of secondary antibodies (goat anti-mouse Alexa488, Thermo; goat anti-rabbit Alexa647, Thermo; Hoechst 33342, 1:2000), and incubated for one hour in the dark. Imaging was performed on a CellInsight CX7 HCS plate reader (Thermo).

### Ingenuity Pathway Analysis

Following differential gene expression analysis of RNA-seq data detailed above, core analyses were accomplished in IPA to identify affected pathways and candidate upstream regulators by comparing gene expression in sham to TAC fibroblasts, and again for TAC to TAC plus JQ1 treatment (FC>+1.5 or FC<-1.5, FDR <0.1). Similarly, differential expression data from the Cuffdiff analysis was used to compare gene expression programs in vehicle to TGF-β treated fibroblasts (log_2_ FC>1 or log_2_<-1, minimum expression 10 fpkm, and FDR <0.1) and again for TGF-β to TGF-β plus JQ1 treated fibroblasts, (log_2_ FC>0.5 or log_2_ FC<-.05, minimum expression 10 fpkm, and FDR<0.1). The top 5 canonical pathways enriched in either analysis was reported with *P*-value in figure format. For Fig. 2c,d, and again in Fig. 3d, genes with expression changes that led to IPA identifying a particular upstream effector molecule (TGF-β, Fig. 2) or pathway (hepatic fibrosis, Fig. 3) were displayed with color indicating strength and direction of expression change (no color if expression change did not meet the above criteria). For illustration purposes in Fig. 2c, the mechanistic network for TGF-β in the analysis was displayed. Then target genes (table provided below the network in IPA) were added to the network. Next, network hubs were deleted (hence some molecules without direct lines from TGF-β). Finally, reduced genes were deleted and induced genes were manually placed to allow visualization. Representation of the same genes to reflect changes from the insult to insult +JQ1 analysis (Fig. 2d and Fig. 3d) was accomplished using the “build” and “grow” options in the Path Designer tool.

## Author contributions

M.S.S., R.A.B, A.S.R. B.Y.E., K.A.K., M.A.C., K.S., M.P.Y.L. performed research and analyzed and interpreted data. M.B.F. and J.Q. provided critical reagents and guidance on experiments. R.A.H. and C.Y.L. analyzed and interpreted ChIP-seq data. M.S.S., J.Q., M.A., C.Y.L., S.M.H. and T.A.M. designed experiments and interpreted data. All authors contributed to the writing of the manuscript.

## Additional information

Supplementary Information accompanies this manuscript.

Competing interests: S.M.H. is an executive and shareholder of Amgen, Inc. and is a shareholder of Tenaya Therapeutics. The other authors declare no competing financial interests.

Accession numbers: The GEO accession number for ChIP-seq data is X, and the accession number for RNA-seq data is X. The publication series accession number is X. Proteomics data are deposited in the PRIDE Archive (https://www.ebi.ac.uk/pride/archive/login). The accession number for the dataset is PXD012482. Username; Password: vqYL216E

## Acknowledgements

We thank C. Danan for RNA-seq library preparation. This work was supported by the NIH (HL116848 to T.A.M. and HL127240 to C.Y.L., S.M.H and T.A.M) and the American Heart Association (16SFRN31400013 to T.A.M). M.S.S. was funded by K01AG056848, 5T32HL007822, and F32HL126354. R.A.B. received funding from the Canadian Institutes of Health Research (FRN-216927), and M.B.F. was supported by an American Heart Association fellowship (19POST34380603). M.P.Y.L. was supported by the NIH (HL127302 and HL141278). M.A. was funded by a Swiss National Science Foundation Postdoctoral Fellowship.

---

**Figure S1. JQ1 suppresses TGF-**β**-induced expression of protein markers of cardiac fibroblast activation**. ARVFs were treated with DMSO vehicle or TGF-β in the absence or presence of JQ1 for 48 hours. Total protein was subjected to LC-MS analysis, as described in the Methods section. (**a**) Label-free quantification of Periostin protein expression. Box: interquartile range; whiskers: min to max, with n=3 plates of cell per condition; Normalized Spectral Abundance Factor (NSAF). (**b**) Principal component analysis of relative protein abundance revealed clear segregation of each treatment group. Each box represents data from an independent plate of ARVFs. (**c**) Heat map of standardized protein values depicts proteins that were increased in expression upon TGF-β stimulation, and impact of JQ1 treatment on this induction (Benjamini & Hochberg adjusted *P*-value <0.1).

**Figure S2. Binding of BRD4 to cardiac fibroblast promoters.** ARVFs were employed for ChIP-seq analysis, as described in Figure 4. Shown is a heat map of BRD4-bound promoters in TGF-β and unstimulated ARVFs covering 1kb upstream and downstream of the transcription start site (TSS); mean reads per million mapped reads per base pair (RPM/bp). Promoters are grouped by gain, loss, or conservation in TGF-β stimulated ARVFs and ranked in order of decreasing BRD4 binding in unstimulated ARVFs in each group.

## References

1. Schuetze, K.B., McKinsey, T.A. & Long, C.S. Targeting cardiac fibroblasts to treat fibrosis of the heart: focus on HDACs. J Mol Cell Cardiol 70, 100–7 (2014).

2. Travers, J.G., Kamal, F.A., Robbins, J., Yutzey, K.E. & Blaxall, B.C. Cardiac Fibrosis: The Fibroblast Awakens. Circ Res 118, 1021–40 (2016).

3. Diez, J. et al. Losartan-dependent regression of myocardial fibrosis is associated with reduction of left ventricular chamber stiffness in hypertensive patients. Circulation 105, 2512–7 (2002).

4. Mohammed, S.F. et al. Coronary microvascular rarefaction and myocardial fibrosis in heart failure with preserved ejection fraction. Circulation 131, 550–9 (2015).

5. Francis Stuart, S.D., De Jesus, N.M., Lindsey, M.L. & Ripplinger, C.M. The crossroads of inflammation, fibrosis, and arrhythmia following myocardial infarction. J Mol Cell Cardiol 91, 114–22 (2016).

6. Moore-Morris, T., Cattaneo, P., Puceat, M. & Evans, S.M. Origins of cardiac fibroblasts. J Mol Cell Cardiol 91, 1–5 (2016).

7. Tallquist, M.D. Cardiac fibroblasts: from origin to injury. Curr Opin Physiol 1, 75–79 (2018).

8. Tallquist, M.D. & Molkentin, J.D. Redefining the identity of cardiac fibroblasts. Nat Rev Cardiol 14, 484–491 (2017).

9. Ivey, M.J. et al. Resident fibroblast expansion during cardiac growth and remodeling. J Mol Cell Cardiol 114, 161–174 (2018).

10. Kanisicak, O. et al. Genetic lineage tracing defines myofibroblast origin and function in the injured heart. Nat Commun 7, 12260 (2016).

11. Kaur, H. et al. Targeted Ablation of Periostin-Expressing Activated Fibroblasts Prevents Adverse Cardiac Remodeling in Mice. Circ Res 118, 1906–17 (2016).

12. Bujak, M. & Frangogiannis, N.G. The role of TGF-beta signaling in myocardial infarction and cardiac remodeling. Cardiovasc Res 74, 184–95 (2007).

13. Davis, J. & Molkentin, J.D. Myofibroblasts: trust your heart and let fate decide. J Mol Cell Cardiol 70, 9–18 (2014).

14. Khalil, H. et al. Fibroblast-specific TGF-beta-Smad2/3 signaling underlies cardiac fibrosis. J Clin Invest 127, 3770–3783 (2017).

15. Molkentin, J.D. et al. Fibroblast-Specific Genetic Manipulation of p38 Mitogen-Activated Protein Kinase In Vivo Reveals Its Central Regulatory Role in Fibrosis. Circulation 136, 549–561 (2017).

16. Stratton, M.S., Koch, K.A. & McKinsey, T.A. p38alpha: A Profibrotic Signaling Nexus. Circulation 136, 562–565 (2017).

17. Davis, J., Burr, A.R., Davis, G.F., Birnbaumer, L. & Molkentin, J.D. A TRPC6-dependent pathway for myofibroblast transdifferentiation and wound healing in vivo. Dev Cell 23, 705–15 (2012).

18. Lighthouse, J.K. & Small, E.M. Transcriptional control of cardiac fibroblast plasticity. J Mol Cell Cardiol 91, 52–60 (2016).

19. Small, E.M. The actin-MRTF-SRF gene regulatory axis and myofibroblast differentiation. J Cardiovasc Transl Res 5, 794–804 (2012).

20. Small, E.M. et al. Myocardin-related transcription factor-a controls myofibroblast activation and fibrosis in response to myocardial infarction. Circ Res 107, 294–304 (2010).

21. Velasquez, L.S. et al. Activation of MRTF-A-dependent gene expression with a small molecule promotes myofibroblast differentiation and wound healing. Proc Natl Acad Sci U S A 110, 16850–5 (2013).

22. Stratton, M.S., Haldar, S.M. & McKinsey, T.A. BRD4 inhibition for the treatment of pathological organ fibrosis. F1000Res 6(2017).

23. Filippakopoulos, P. et al. Selective inhibition of BET bromodomains. Nature 468, 1067–73 (2010).

24. Anand, P. et al. BET bromodomains mediate transcriptional pause release in heart failure. Cell 154, 569–82 (2013).

25. Duan, Q. et al. BET bromodomain inhibition suppresses innate inflammatory and profibrotic transcriptional networks in heart failure. Sci Transl Med 9(2017).

26. Spiltoir, J.I. et al. BET acetyl-lysine binding proteins control pathological cardiac hypertrophy. J Mol Cell Cardiol 63, 175–9 (2013).

27. Arora, P.D. & McCulloch, C.A. Dependence of collagen remodelling on alpha-smooth muscle actin expression by fibroblasts. J Cell Physiol 159, 161–75 (1994).

28. Bell, E., Ivarsson, B. & Merrill, C. Production of a tissue-like structure by contraction of collagen lattices by human fibroblasts of different proliferative potential in vitro. Proc Natl Acad Sci U S A 76, 1274–8 (1979).

29. Montesano, R. & Orci, L. Transforming growth factor beta stimulates collagen-matrix contraction by fibroblasts: implications for wound healing. Proc Natl Acad Sci U S A 85, 4894–7 (1988).

30. Brown, J.D. et al. NF-kappaB directs dynamic super enhancer formation in inflammation and atherogenesis. Mol Cell 56, 219–231 (2014).

31. Chapuy, B. et al. Discovery and characterization of super-enhancer-associated dependencies in diffuse large B cell lymphoma. Cancer Cell 24, 777–90 (2013).

32. Loven, J. et al. Selective inhibition of tumor oncogenes by disruption of super-enhancers. Cell 153, 320–34 (2013).

33. Stratton, M.S. et al. Signal-Dependent Recruitment of BRD4 to Cardiomyocyte Super-Enhancers Is Suppressed by a MicroRNA. Cell Rep 16, 1366–1378 (2016).

34. Bisgrove, D.A., Mahmoudi, T., Henklein, P. & Verdin, E. Conserved P-TEFb-interacting domain of BRD4 inhibits HIV transcription. Proc Natl Acad Sci U S A 104, 13690–5 (2007).

35. Jang, M.K. et al. The bromodomain protein Brd4 is a positive regulatory component of P-TEFb and stimulates RNA polymerase II-dependent transcription. Mol Cell 19, 523–34 (2005).

36. Yang, Z. et al. Recruitment of P-TEFb for stimulation of transcriptional elongation by the bromodomain protein Brd4. Mol Cell 19, 535–45 (2005).

37. Roe, J.S., Mercan, F., Rivera, K., Pappin, D.J. & Vakoc, C.R. BET Bromodomain Inhibition Suppresses the Function of Hematopoietic Transcription Factors in Acute Myeloid Leukemia. Mol Cell 58, 1028–39 (2015).

38. Arabacilar, P. & Marber, M. The case for inhibiting p38 mitogen-activated protein kinase in heart failure. Front Pharmacol 6, 102 (2015).

39. Bennetts, J.S. et al. Evolutionary conservation and murine embryonic expression of the gene encoding the SERTA domain-containing protein CDCA4 (HEPP). Gene 374, 153–65 (2006).

40. Calgaro, S., Boube, M., Cribbs, D.L. & Bourbon, H.M. The Drosophila gene taranis encodes a novel trithorax group member potentially linked to the cell cycle regulatory apparatus. Genetics 160, 547–60 (2002).

41. Watanabe-Fukunaga, R., Iida, S., Shimizu, Y., Nagata, S. & Fukunaga, R. SEI family of nuclear factors regulates p53-dependent transcriptional activation. Genes Cells 10, 851–60 (2005).

42. Hsu, S.I. et al. TRIP-Br: a novel family of PHD zinc finger-and bromodomain-interacting proteins that regulate the transcriptional activity of E2F-1/DP-1. EMBO J 20, 2273–85 (2001).

43. Sugimoto, M. et al. Regulation of CDK4 activity by a novel CDK4-binding protein, p34(SEI-1). Genes Dev 13, 3027–33 (1999).

44. Kusano, S., Shiimura, Y. & Eizuru, Y. I-mfa domain proteins specifically interact with SERTA domain proteins and repress their transactivating functions. Biochimie 93, 1555–64 (2011).

45. Peng, Y., Zhao, S., Song, L., Wang, M. & Jiao, K. Sertad1 encodes a novel transcriptional co-activator of SMAD1 in mouse embryonic hearts. Biochem Biophys Res Commun 441, 751–6 (2013).

46. Song, S. et al. Inhibition of BRD4 attenuates transverse aortic constriction-and TGF-beta-induced endothelial-mesenchymal transition and cardiac fibrosis. J Mol Cell Cardiol 127, 83–96 (2018).

47. Tang, X. et al. Assessment of Brd4 inhibition in idiopathic pulmonary fibrosis lung fibroblasts and in vivo models of lung fibrosis. Am J Pathol 183, 470–9 (2013).

48. Tang, X. et al. BET bromodomain proteins mediate downstream signaling events following growth factor stimulation in human lung fibroblasts and are involved in bleomycin-induced pulmonary fibrosis. Mol Pharmacol 83, 283–93 (2013).

49. Tian, B. et al. BRD4 mediates NF-kappaB-dependent epithelial-mesenchymal transition and pulmonary fibrosis via transcriptional elongation. Am J Physiol Lung Cell Mol Physiol 311, L1183–L1201 (2016).

50. Zhou, B. et al. Brd4 inhibition attenuates unilateral ureteral obstruction-induced fibrosis by blocking TGF-beta-mediated Nox4 expression. Redox Biol 11, 390–402 (2017).

51. Ding, N. et al. BRD4 is a novel therapeutic target for liver fibrosis. Proc Natl Acad Sci U S A 112, 15713–8 (2015).

52. Kumar, K. et al. BET inhibitors block pancreatic stellate cell collagen I production and attenuate fibrosis in vivo. JCI Insight 2, e88032 (2017).

53. Andrieu, G., Belkina, A.C. & Denis, G.V. Clinical trials for BET inhibitors run ahead of the science. Drug Discov Today Technol 19, 45–50 (2016).

54. Doroshow, D.B., Eder, J.P. & LoRusso, P.M. BET inhibitors: a novel epigenetic approach. Ann Oncol 28, 1776–1787 (2017).

55. Bagchi, R.A. et al. The transcription factor scleraxis is a critical regulator of cardiac fibroblast phenotype. BMC Biol 14, 21 (2016).

56. Lighthouse, J.K. et al. Exercise promotes a cardioprotective gene program in resident cardiac fibroblasts. JCI Insight 4(2019).

57. Adusumilli, R. & Mallick, P. Data Conversion with ProteoWizard msConvert. Methods Mol Biol 1550, 339–368 (2017).

58. Eng, J.K. et al. A deeper look into Comet--implementation and features. J Am Soc Mass Spectrom 26, 1865–74 (2015).

59. UniProt, C. UniProt: a hub for protein information. Nucleic Acids Res 43, D204–12 (2015).

60. The, M., MacCoss, M.J., Noble, W.S. & Kall, L. Fast and Accurate Protein False Discovery Rates on Large-Scale Proteomics Data Sets with Percolator 3.0. J Am Soc Mass Spectrom 27, 1719–1727 (2016).

61. Ritchie, M.E. et al. limma powers differential expression analyses for RNA-sequencing and microarray studies. Nucleic Acids Res 43, e47 (2015).

62. Huber, W. et al. Orchestrating high-throughput genomic analysis with Bioconductor. Nat Methods 12, 115–21 (2015).

63. Dobin, A. et al. STAR: ultrafast universal RNA-seq aligner. Bioinformatics 29, 15–21 (2013).

64. Team, R.C. R: A Language and Environment for Statistical Computing. https://www.R-project.org/ (2017).

65. Robinson, M.D., McCarthy, D.J. & Smyth, G.K. edgeR: a Bioconductor package for differential expression analysis of digital gene expression data. Bioinformatics 26, 139–40 (2010).

66. Trapnell, C. et al. Transcript assembly and quantification by RNA-Seq reveals unannotated transcripts and isoform switching during cell differentiation. Nat Biotechnol 28, 511–5 (2010).

67. Langmead, B. & Salzberg, S.L. Fast gapped-read alignment with Bowtie 2. Nat Methods 9, 357–9 (2012).

68. Zhang, Y. et al. Model-based analysis of ChIP-Seq (MACS). Genome Biol 9, R137 (2008).

