## Supplement for "Dynamic chromatin targeting of BRD4 stimulates cardiac fibroblast activation"

### sFigure 1

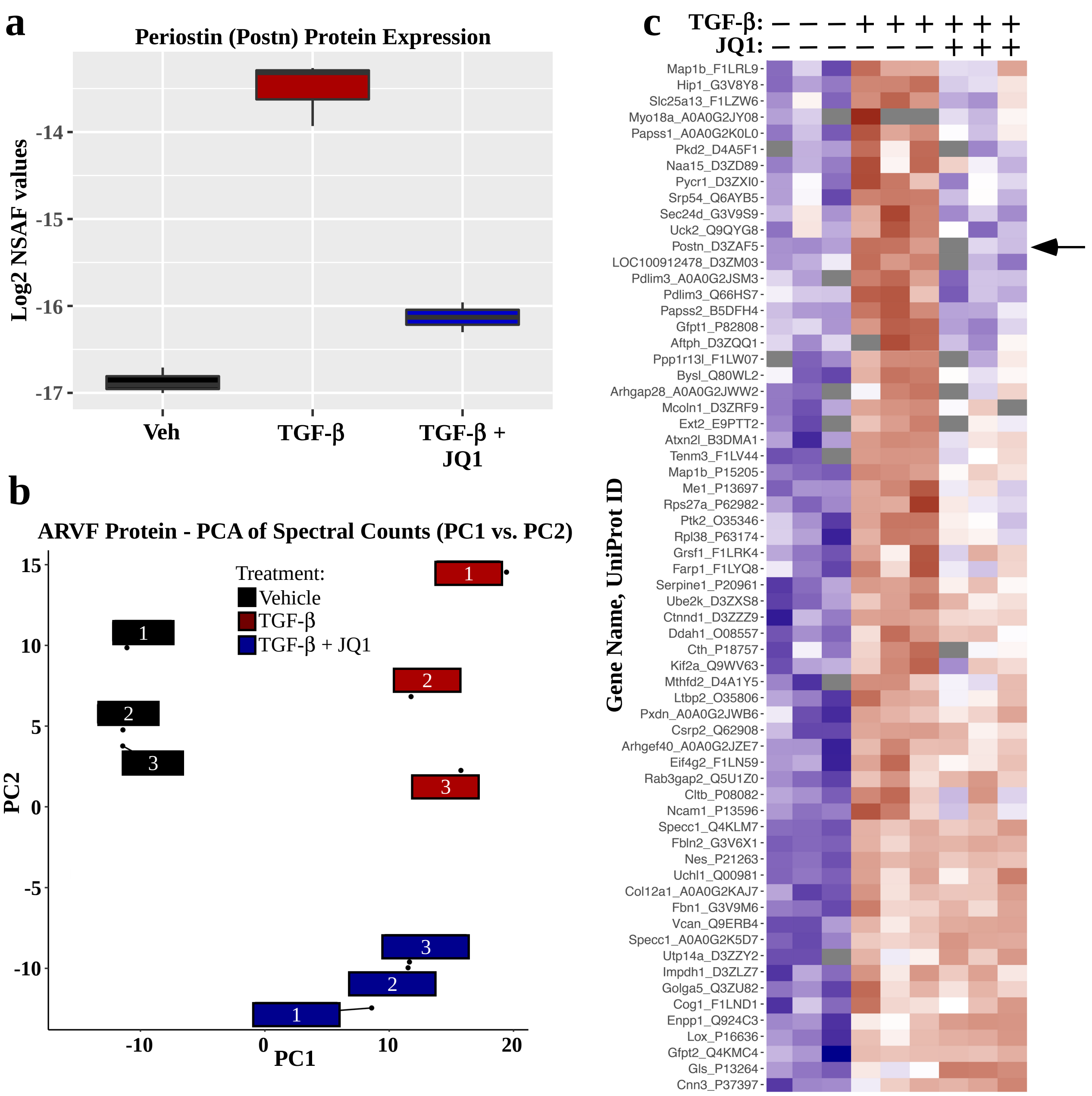
a
Periostin (Postn) Protein Expression
Log2 NSAF values
-14
-15
-16
-17
Veh
TGF-β
TGF-β + JQ1
b
ARVF Protein - PCA of Spectral Counts (PC1 vs. PC2)
Treatment:
Vehicle
TGF-β
TGF-β + JQ1
PC2
15
10
5
0
-5
-10
PC1
-10
0
10
20
c
TGF-β: − − − + + + + + +
JQ1: − − − − − − + + +
Gene Name, UniProt ID
Map1b_F1LRL9
Hip1_G3V8Y8
Slc25a13_F1LZW6
Myo18a_A0A0G2JY08
Papss1_A0A0G2K0L0
Pkd2_D4A5F1
Naa15_D3ZD89
Pycr1_D3ZXI0
Srp54_Q6AYB5
Sec24d_G3V9S9
Uck2_Q9QYG8
Postn_D3ZAF5
LOC100912478_D3ZM03
Pdlim3_A0A0G2JSM3
Pdlim3_Q66HS7
Papss2_B5DFH4
Gfpt1_P82808
Aftph_D3ZQQ1
Ppp1r13l_F1LW07
Bysl_Q80WL2
Arhgap28_A0A0G2JWW2
Mcoln1_D3ZRF9
Ext2_E9PTT2
Atxn2l_B3DMA1
Tenm3_F1LV44
Map1b_P15205
Me1_P13697
Rps27a_P62982
Ptk2_O35346
Rpl38_P63174
Grsf1_F1LRK4
Farp1_F1LYQ8
Serpine1_P20961
Ube2k_D3ZXS8
Ctnnd1_D3ZZZ9
Ddah1_O08557
Cth_P18757
Kif2a_Q9WV63
Mthfd2_D4A1Y5
Ltbp2_O35806
Pxdn_A0A0G2JWB6
Csrp2_Q62908
Arhgef40_A0A0G2JZE7
Eif4g2_F1LN59
Rab3gap2_Q5U1Z0
Cltb_P08082
Ncam1_P13596
Specc1_Q4KLM7
Fbln2_G3V6X1
Nes_P21263
Uchl1_Q00981
Col12a1_A0A0G2KAJ7
Fbn1_G3V9M6
Vcan_Q9ERB4
Specc1_A0A0G2K5D7
Utp14a_D3ZZY2
Impdh1_D3ZLZ7
Golga5_Q3ZU82
Cog1_F1LND1
Enpp1_Q924C3
Lox_P16636
Gfpt2_Q4KMC4
Gls_P13264
Cnn3_P37397

### sFigure 2

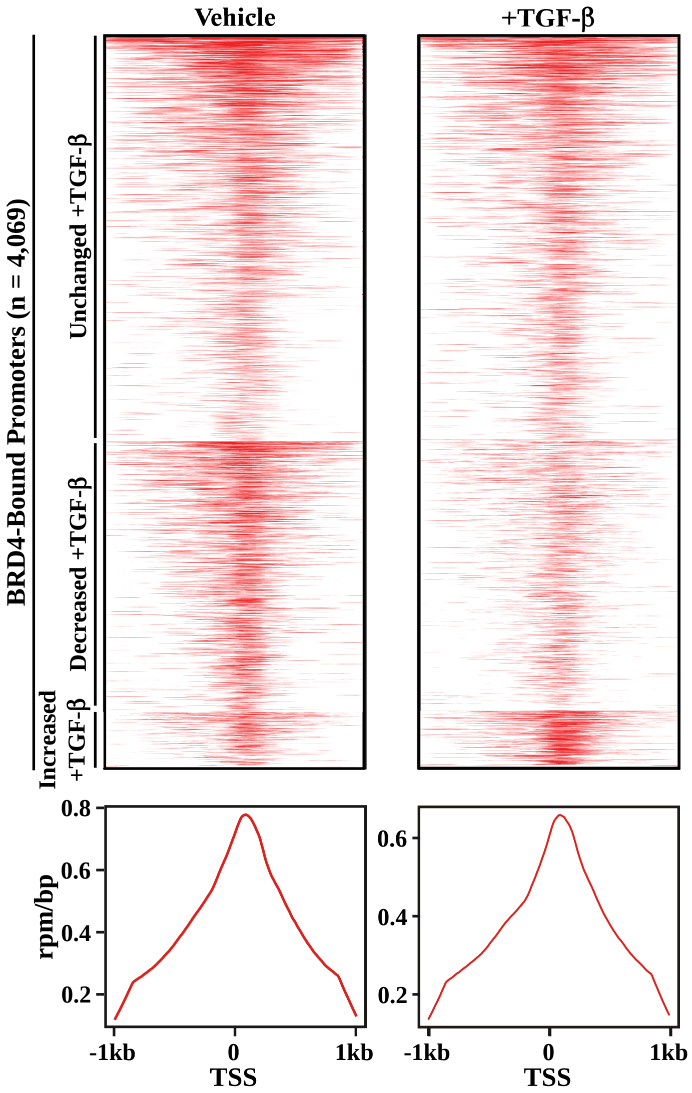
Vehicle
+TGF-β
BRD4-Bound Promoters (n = 4,069)
Unchanged +TGF-β
Decreased +TGF-β
Increased +TGF-β
rpm/bp
0.8
0.6
0.4
0.2
-1kb
0
1kb
TSS

**Supplemental Table 1: Primers for qPCR**

| Target | 5' oligo | 3' oligo |
|---|---|---|
| *Brd2* | AGCACCAGGGAAAAGGATTC | GATGCTTCCACAGAGCCTTC |
| *Brd3* | CTCCCCCACCTGAGGTCT | AGGCAGGTTCAGCTTGATTG |
| *Brd4* | CCCTGGGACAAGATTGAGAA | CTGCCTCTTGGGCTTGTTAG |
| *B2M* | TCACACCCACCGAGACCGATGT | TCTCGGTCCCAGGTGACGGTTT |
| *Sertad4* | GGTAAGTTCTCCCACTTTCTCC | TGCCCAAACGATCTACTCATAC |
| *αSMA* | GGAGATGGCGTGACTCACAA | CGCTCAGCAGTAGTCACGAA |
| *Periostin* | TCGTGGAACCAAAAATTAAAGTC | CTTCGTCATTGCAGGTCCTT |

**Supplemental Table 2: Primers for ChIP-qPCR**

| Target | 5' oligo | 3' oligo |
|---|---|---|
| *Sertad4 Enh/SE 1* | CAAATGCTGGGAATCTGAACTG | TTAGGCCCTTCTGTCCTATCT |
| *Sertad4 Enh/SE 2* | CCACTTCAGAGACAAGCTCTTT | CACAGACTAGGTTGAGCATTCC |
| *Sertad4 Enh 3* | CACTACCGCCACTTACCTTTC | GGCGACTTCTCTGAAACCAATA |
| *Sertad4 Enh 4* | ACAGTGCCAGAGTCCAAAC | CTCTGATGGTGCTGACCTATTT |
| *Sertad4 Enh 5* | TTCGGGCATCCGTTCAAA | TGCTTGGAGCGCTTTCTTA |
| *Sertad4 Enh 6* | CAGGGCTGGAAGGAAAGTAAA | CGAGGTTTATGGCGGGATTAG |
| *Sertad4 TSS* | CAGTGCCTCTCCTGAAGAATAA | GGCGTGTGTTTCCCATAGA |
